# Caloric restriction prevents inflammation and insulin dysfunction in middle-aged ovariectomized mice

**DOI:** 10.1101/2023.02.09.527836

**Authors:** Leticia Roberta Leme Sapatini, Bruno Calsa, Lais Jorge Marim, Júlia Venturini Helaehil, Gabriela Bortolança Chiarotto, Maria Esméria Corezola do Amaral

**Author notes:** **Corresponding author:** Maria Esméria Corezola do Amaral, Graduate Program in Biomedical Science, University Center of Hermínio Ometto Foundation, Av. Maximiliano Barutto nº 500, Jardim Universitário, 13607-339, Araras, SP, Brazil. Co-first authors: Leticia Sapatini and Bruno Calsa.

## Abstract

Loss of ovarian function is associated with increased visceral fat. In this study, we aimed to study the effects of caloric restriction (CR) on metabolism in ovariectomized mice. Female, 8–12-month-old mice were divided into three groups: OVX (ovariectomized mice), OVXR (40% CR) and Sham. CR increased insulin sensitivity and glucose tolerance. AMPK phosphorylation was observed in the liver of OVXR mice. CR also increased hepatic cholesterol and triglyceride levels. The reductions in the level of TBARS in the serum and liver and of H2O2 in the liver of OVXR mice suggested alterations in the redox state of the liver. Although expression of catalase protein was reduced by CR, expression of superoxide dismutase was not altered by CR. Although interleukin IL-6 and IL-10 levels in OVXR mice were similar to those in Sham mice, macrophage infiltration was reduced in OVXR mice. OVXR mice had increased sirtuin1 levels and decreased sirtuin3 levels in the liver. In conclusion, CR improved the condition of ovariectomized mice by reducing adiposity and increasing insulin sensitivity and glucose tolerance through a mechanism that may involve AMPK.

## Introduction

Numerous studies have shown that menopause is associated with unfavorable changes in body composition, such as increased abdominal fat deposition, and poor overall health [1, 2]. The deleterious effects of obesity are diverse and include various metabolic diseases, such as diabetes mellitus, and cardiovascular diseases, like coronary heart disease, heart attack, stroke, and hypertension, which adversely impact the quality of life and carry an increased risk of premature death [3-6]. However, the relationship between menopause and obesity is complex. Sex hormones fluctuate with reproductive changes, such as menarche, pregnancy, and menopause, which may play an important role in the expansion of adipose tissue [1]. Hormones impact adipose tissue, and estrogens regulate glucose homeostasis and insulin sensitivity in several ways, by altering insulin production and secretion [7, 8], insulin-mediated glucose uptake in peripheral tissues, mitochondrial function [9], and the expression of inflammatory markers [10]. Given the apparent interaction between low estrogen concentrations and obesity in altering glucose and lipid homeostasis, increasing attention has been given to developing dietary strategies that preserve homeostasis during menopause.

Caloric restriction (CR) is one strategy that can reduce body weight, total body fat, and visceral fat, similar to exercise. CR is a non-pharmacological dietary intervention in which the average daily calorie intake is reduced without malnutrition or deprivation of essential nutrients. Animals on a calorie-restricted diet showed a significant (50%) increase in life expectancy [11]. Animal studies showed favorable effects of short-term CR on cardiometabolic changes, liver changes, and cellular senescence [12, 13]. In humans, experimental and observational studies have revealed that CR can reduce the incidence of cardiovascular disease, diabetes, dementia, frailty, and cancer [14, 15]. These observed benefits are related to CR-mediated activation of regulatory proteins in metabolic pathways, such as AMP-activated protein kinase (AMPK) and sirtuins (SIRTs). AMPK is a key sensor and regulator of glucose metabolism that activates SIRT1 deacetylase in response to changes in the redox state (NAD+/NADH and AMP/ATP ratios) of the cell [16]. Sirtuins are a group of highly conserved NAD+-dependent histones and protein deacetylases and/or ADP-ribosyl transferases that play important roles in various biological processes [17, 18, 8, 19]. SIRT1 functions as a metabolic/energy sensor that modulates the activity and/or gene expression of several crucial transcription factors and transcription coactivators involved in metabolic homeostasis [20, 18, 21]. Reduced levels of NAD+ and SIRT1 activity have been detected under energy-overloaded metabolic profiles, such as a hyperlipidic diet [22, 23].

In contrast, CR increased NAD+ levels and induced activation of SIRT1 [22, 23]. SIRT1 functions as an essential regulator of hepatic lipid metabolism through SREBP-1c/ChREBP-dependent control of lipogenesis by inhibiting lipid synthesis and increasing PPARα/PGC-1α-dependent β-fatty acid oxidation [24]. SIRT1 regulates oxidative stress by increasing the transcriptional activity of antioxidant enzymes, such as superoxide dismutase (SOD), catalase (CAT), and glutathione peroxidase (GPX) [24]. Activation of SIRT1 reduces nitric oxide synthase and lipid peroxidation products, which has an anti-oxidative stress effect in the liver [24]. SIRT3 is a mitochondrial protein that integrates cellular energy metabolism and plays an important role in preventing metabolic syndrome [25]. SIRT3 knockout mice showed characteristics of reduced mitochondrial metabolism, such as reduced fatty acid catabolism [25]. SIRT3 expression was reduced in male mice fed a high-fat diet [25, 26]. However, the consistency and reproducibility of results across investigations are varied, and key questions about the timing, duration, modality, and adverse effects of CR, as well as the influence of genotype and diet composition, remain [27]. Given the low adherence of humans to difficult-to-follow diets, research on dietary interventions should be conducted. This study examined the impact of 40% CR on the liver of middle-aged mice who underwent ovariectomy. CR was expected to have a positive effect on metabolism. The results showed that CR was associated with improvements in both metabolism, mainly glycemic metabolism, and inflammatory markers.

## Materials and methods

### Experimental animals and model establishment

The project was approved by the Committee on Ethics in Animal Use (CEUA) under protocol #009/2020. Eighteen C57BL/6J mice (8–12 months old, weighing ∼20 g) were housed in individual cages at a constant temperature (22 °C) under a 12 h light/dark cycle with water and commercial chow *ad libitum*. For the ovariectomy, the mice were anesthetized by intraperitoneal injection with ketamine (50 mg/kg) and xylazine (5 mg/kg). After shaving the inguinal region, an abdominal incision was made, and the ovaries were located and removed. After surgery, the animals were placed in a heated chamber to avoid hypothermia. The animals in the control group (Sham) underwent the same procedure except the ovaries were not removed. At 16 weeks after surgery, the ovariectomy model mice were separated into two groups: the OVX (ovariectomized) and OVXR (ovariectomized plus 40% CR) groups. Sham and OVX mice were fed a commercial isocaloric diet *ad libitum*, and OVXR mice received 60% of the chow consumed by the OVX mice for 30 days, corresponding to 40% CR. All mice had free access to water throughout the experiment. The body weights of the animals were measured weekly throughout the experiment.

### Intraperitoneal glucose tolerance test

The intraperitoneal glucose tolerance test (ipGTT) was administered 5 days before euthanasia after a 6 h fast. For the test, glucose (2 g/kg body weight) was injected intraperitoneally. Blood samples were collected before glucose overload (time zero), and at 15, 30, 60, and 120 min after glucose infusion. Blood glucose was determined using reagent strips and an Abbott® glucometer as previously described [28].

### Intraperitoneal insulin tolerance test

The intraperitoneal insulin tolerance test (ipITT) was administered 3 days before euthanasia after a 6 h fast. Insulin (regular crystalline, 0.75 U/kg body weight) was administered intraperitoneally. Blood samples were collected before insulin injection (time zero) and at 5, 10, 15, 20, 25, and 30 min after injection. Blood glucose was determined using an Abbott glucometer and reagent strips. The rate constant for glucose disappearance (Kitt) was calculated using the formula In2/t1/2. The half-life (t1/2) of serum glucose was calculated from the slope of the minimum regression curve in the linear phase of plasma glucose decline [29].

### Homeostasis model assessments

Homeostasis model assessment of insulin resistance (HOMA-IR), insulin sensitivity (HOMA%S), and β-cell function (HOMA%β) were calculated using the HOMA calculator from the University of Oxford (www.dtu.ox.ac.uk/homacalculator).

### Fat pad and Lee indexes

The perirenal adipose tissue weight and body weight were assessed, and the adipose tissue index was calculated by dividing adipose tissue weight by body weight. The Lee index was calculated using the following equation: 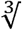body weight (g)÷nose to anus length (cm)..

### Biochemical assessments

Serum glucose, triglycerides, cholesterol, HDL, and total protein were determined using commercial kits (Laborlab, Guarulhos, SP, Brazil). LDL was calculated using the Friedewald equation, and VLDL was calculated by dividing triglycerides by 5. Insulin serum was measured using ELISA (Rat/Mouse Insulin ELISA, #EZRMI-13K, Millipore St. Charles, MO, USA)). Hepatic glycogen content was measured using a colorimetric method [30]. Total liver lipids were extracted according to the method of Folch et al. [31].

### Preparation of liver homogenates for oxidative stress biomarker analysis

Liver samples (∼50 mg) were homogenized in Krebs buffer (118 mM NaCl, 25 mM NaHCO_3_, 1.2 mM KH_2_PO_4_, 4.7 mM KCl, 1.2 mM MgSO_4_, 1.25 mM CaCl_2_, 10 mM glucose, and 10 mM HEPES, pH 7.4). The obtained tissue homogenate was used for the determination of 2-thiobarbituric acid reactive substances (TBARS) and hydrogen peroxide (H_2_O_2_) levels. Total protein was quantified using the Biuret method, with bovine serum albumin as a standard, and measured at 540 nm.

### Determination of hepatic H_2_O_2_

H_2_O_2_ levels in hepatic tissue were determined using the Amplex Red method. Briefly, 20 µL of homogenate was incubated with horseradish peroxidase (HRP, 1 U/mL) and Amplex Red (50 μM) at 37 °C for 30 min. Then, the absorbance was read at 560 nm. The H_2_O_2_ concentration was calculated by referring to a standard curve of known concentrations and normalized to the protein concentration.

### Hepatic and systemic TBARS

Serum and liver TBARS contents were determined as described previously [32]. Briefly, 50 µL of serum or liver homogenate was mixed with 10 µL of 0.1 M Butylated Hydroxytoluene, 400 µL of 1% 2-thiobarbituric acid (TBA; Merck, St. Charles, MO, USA), and 200 µL of 20% phosphoric acid and heated at 100 °C for 15 min. After cooling, 1.5 mL of 1-butanol was added, and the optical density of the supernatant was determined at 532 nm.

### Determination of hepatic n-acetylglucosaminidase

n-Acetylglucosaminidase (NAG) activity was measured to determine liver macrophagic infiltration. An aliquot (3.0 μL) of liver homogenate (0.08 M NaPO_4_) was mixed with 30 μL of p-nitrophenyl-2-acetamide-β-D-glucopyranoside (Sigma-Aldrich) and diluted in 50 μL of 50 mM citrate buffer. Finally, 50 μL of 0.2 M glycine was added, and the absorbance was read at 405 nm.

### Histological analysis of liver and adipose tissue

Liver and adipose tissues were fixed with 10% formalin and dehydrated. Then, small fragments of the tissues were embedded in paraffin (Paraplast; Sigma, St. Louis, MO, USA) using standard procedures. Slices (5 μm thick) were stained with hematoxylin-eosin (HE), and images were acquired using a microscope (Leica DM 2000, Germany) with LAS v.4.1 software. For each mouse, 40 images were obtained at 400× magnification. The area (mm^2^) containing adipocytes in each image was quantified using the ImageJ software (National Institutes of Health, Bethesda, Maryland, USA).

### Immunoblotting

Liver tissue fragments were homogenized in protein extraction buffer (10 mM EDTA, 100 mM Tris-HCL, 10 mM sodium pyrophosphate, 100 mM sodium fluoride, 100 mM sodium orthovanadate, 10 mM phenylmethylsulfonyl fluoride, and 0.1 mg/mL aprotinin). Protein content was determined according to the Biuret method. The samples were mixed with Laemmli buffer (0.1% bromophenol blue, 1 M sodium phosphate, 50% glycerol, and 10% SDS). Total protein (50 μg) was separated by electrophoresis. Then, the separated proteins were transferred to nitrocellulose membranes. After transfer, the membranes were incubated at 4 °C, with shaking, overnight with the following primary antibodies: IL-6, IL-10, TGF-β1, SIRT3, pIR, and IR (Santa Cruz, Dallas, USA; 1:250), and SIRT1, pAMPK, and AMPK (Cell Signaling Danvers, Massachusetts, USA; 1:1000). The membranes were then incubated with an HRP-secondary antibody (Santa Cruz; Dallas, USA; 1:10,000), with shaking, followed by development with chemiluminescent reagents (SuperSignal West Pico PLUS, Thermo Scientific, Rockford, IL, USA) and bands intensities were determined by densitometry scanning using the ImageJ software (National Institutes of Health, Bethesda, Maryland, USA)).

### Statistical analysis

Data are mean ± standard error of the mean. All data were analyzed using the Shapiro– Wilk Normality test. For comparisons between experimental groups, one-way ANOVA and Tukey’s post-test were used. All statistical and graphical analyses were performed using GraphPad Prism software (version 8.0). Significance was set at p <0.05.

## Results

### Ovariectomy and caloric restriction characterization

The timeline of the experiment is shown in Figure 1A. After ovariectomy, the animals were observed for 4 months. The animals in the OVXR group remained on a restricted diet for the first month. The body weights of the animals were monitored throughout the experiment and are presented in Figure 1B, which shows evident weight gain in the OVX group compared to in the Sham group and marked weight loss during CR in the OVXR group compared to in the Sham and OVX groups. The Lee index of the mice was lower in the OVXR group than in the Sham and OVX groups (Figure 1C). Analysis of the uterine weight of the mice showed evident uterine atrophy following ovariectomy (Figure 1D).

**Fig. 1.**
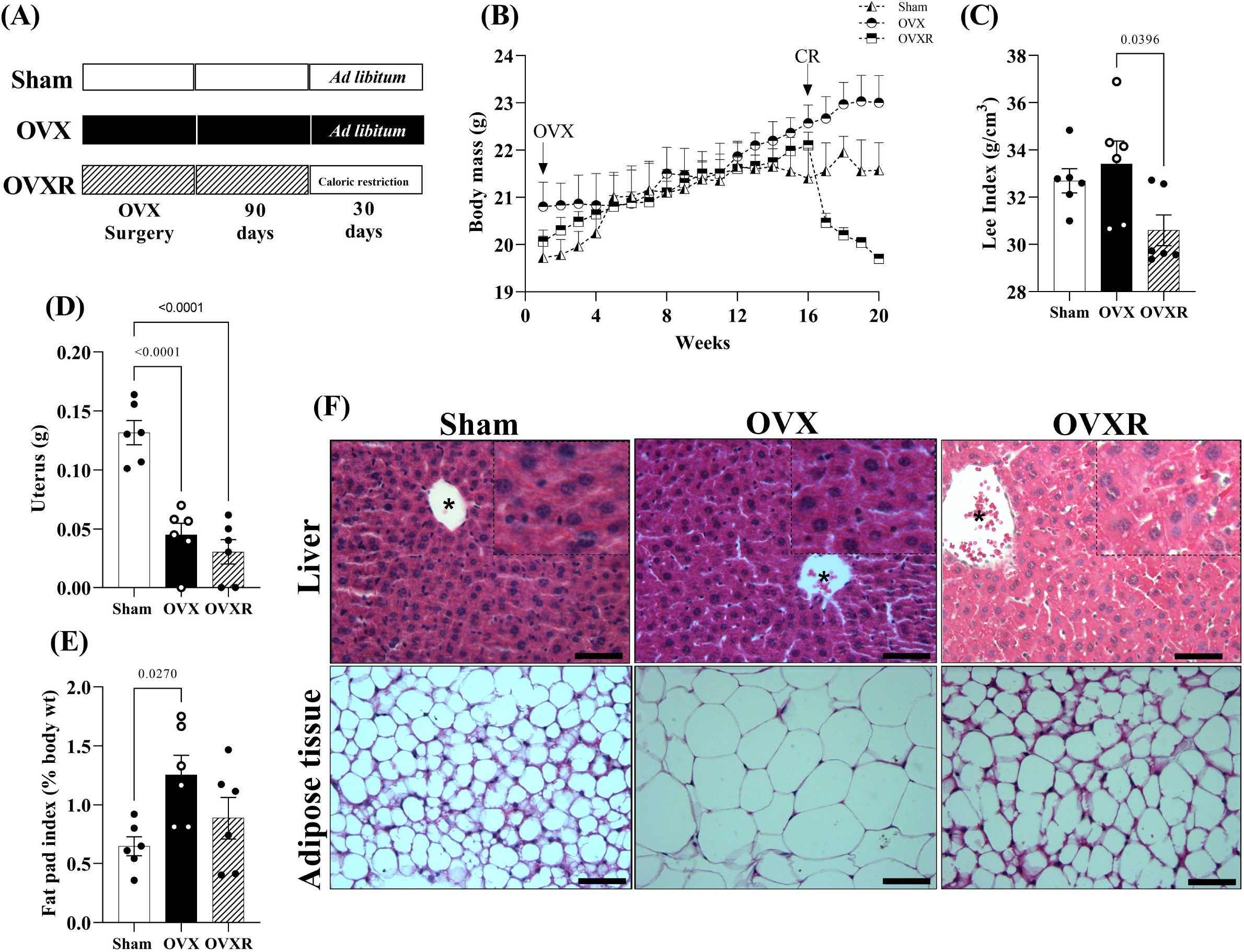
Metabolic effects of caloric restriction on ovariectomized mice. (A) Experiment timeline. (B) Body mass, (C) Lee index, (D) uterine weight, (E) Fat pad index, and (F) histological examination of the liver and adipose tissue in sham operated (Sham), ovariectomized (OVX), and ovariectomized + 40% caloric restriction (CR) (OVXR) mouse groups. Data are mean ± SEM and were analyzed using one-way ANOVA followed by post-hoc Tukey test. The numbers of mice (n) and p-values are indicated on each graph.

Mice in the OVXR group had lower perirenal adipose tissue weight than mice in the OVX group (Figure 1E); however, the difference was not statistically different. OVX mice showed adipose tissue weight gain when compared to mice in the other groups. HE staining of liver and adipose tissue (Figure 1F) showed slight steatosis in OVXR group mice when compared to the Sham and OVX group mice. There was an evident reduction in adipocyte diameter in OVXR mice when compared to that in OVX mice.

### Effects on glucose, insulin, and lipid metabolism

The GTT and ITT results (Figures 2A and 2B, respectively) showed a reduced area under the curve for OVXR mice (Figure 2C) compared to that for OVX mice and an increased Kitt for OVXR mice compared to that for OVX mice (Figure 2D). These results suggest greater glucose tolerance and increased insulin sensitivity of OVXR mice than of OVX mice. The HOMA-IR (Figure 2E), HOMA%S (Figure 2F), and HOMA%B (Figure 2G) indexes indicated increased insulin sensitivity and beta cell preservation in the OVXR mice compared to in the OVX mice. Consistent with these results, liver glycogen levels were similar in OVXR and Sham mice but lower in OVX than in OVXR mice (Table 1). No difference in the liver index was observed among the groups. The reduced blood glucose levels (Figure 3F) and insulinemia (Figure 3G) in the OVXR group compared to that in the Sham and OVX groups suggest improvement in glycemic homeostasis under CR. Similar serum protein concentrations (Figure 3H) in all groups indicate good nutritional status.

**Fig. 2.**
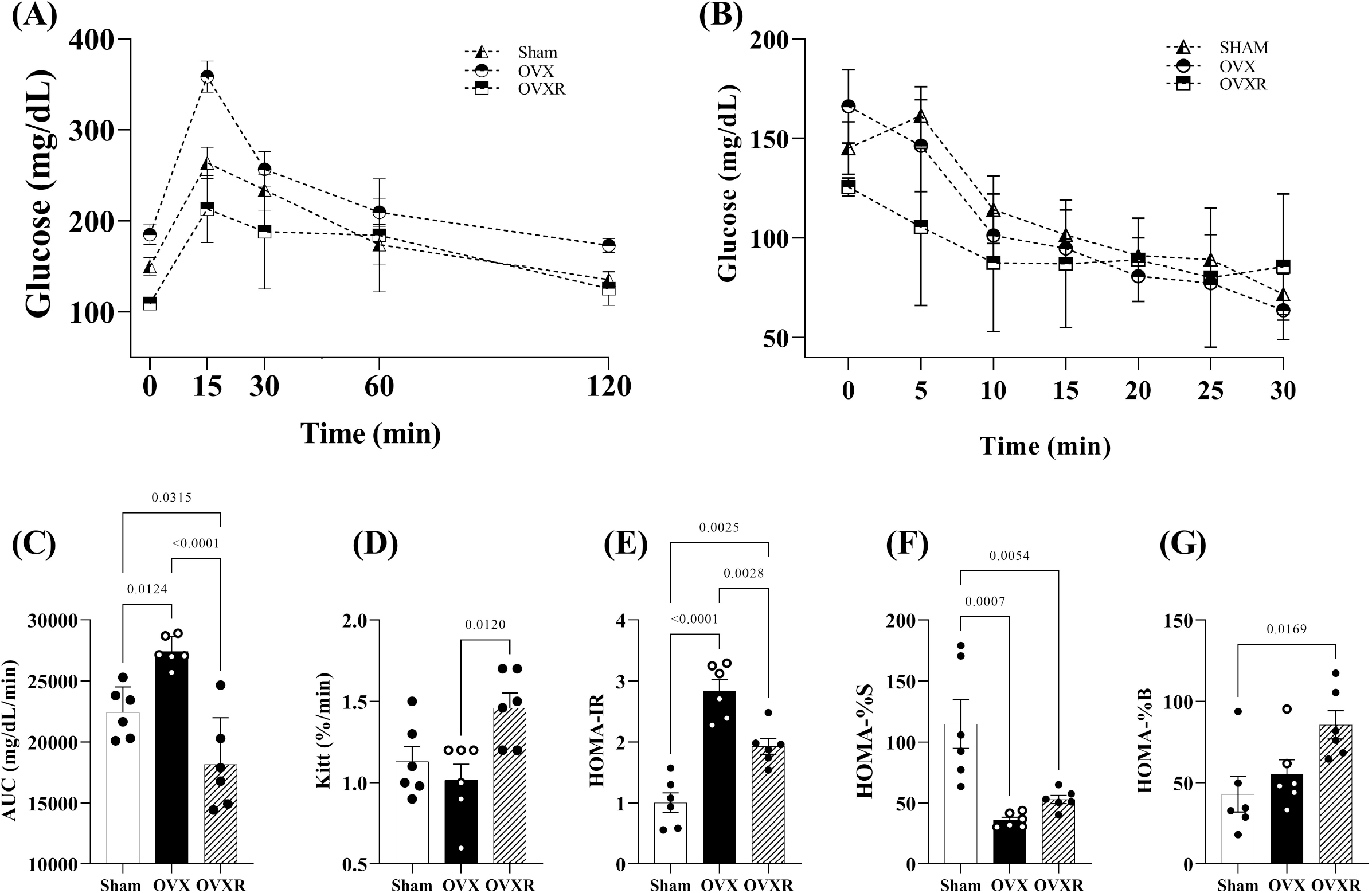
Biochemical analyses of glycemic homeostasis. (A) Intraperitoneal glucose tolerance test (ipGTT); (B) intraperitoneal insulin tolerance test (ipITT); (C) area under the curve (AUC) of the glucose tolerance test; (D) rate constant for glucose decay (Kitt) during the ITT test; and (E) HOMA-IR, (F) HOMA%S, and (G) HOMA%B indexes. Data are mean ± SEM and were analyzed by one-way ANOVA followed by post-hoc Tukey test. The numbers of mice (n) and p-values are indicated on each graph.

**Fig. 3.**
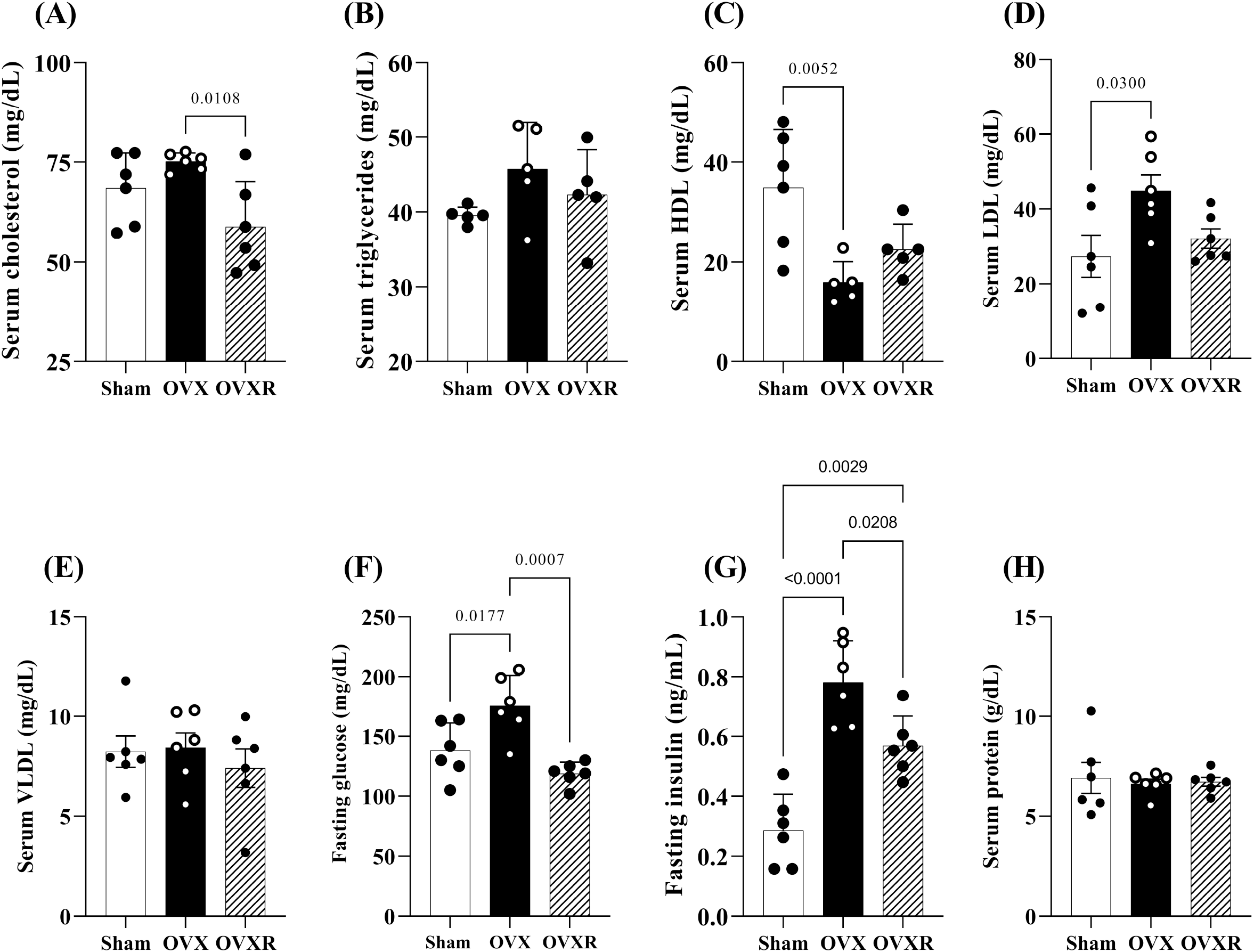
Biochemical analyses of lipid metabolism. Measurements of (A) serum cholesterol, (B) serum triglycerides, (C) serum HDL, (D) serum LDL, (E) serum VLDL, (F) fasting glucose, (G) fasting insulin, and (H) serum protein. Data are mean ± SEM and were analyzed by one-way ANOVA followed by post-hoc Tukey test. The numbers of mice (n) and p-values are indicated on each graph.

**Table 1:** Hepatic and adipose characteristics.

| Parameters | Sham | OVX | OVXR | Statistical difference |
| --- | --- | --- | --- | --- |
| **Hepatic Chol**<br>**(mg/100g tissue)** | 0.001435±0.0001 | 0.001637±0.0001 | 0.001796±0.0003 | None |
| **Hepatic TG**<br>**(mg/100g tissue)** | 0.0005738±0.0001 | 0.0006981±0.0002 | 0.001502±0.0003<sup>a,b</sup> | a: p=0.003 vs Sham<br>b: p=0.008 vs OVX |
| **Hepatic Glycogen**<br>**(mg/100g tissue)** | 0.8490±0.144 | 0.2902±0.059<sup>a,c</sup> | 1.164±0.131 | a: p=0.012 vs Sham<br>c: p=0.0001 vs OVXR |
| **Liver tissue index\*** | 4.660±0.2420 | 4.454±0.1980 | 4.930±0.4340 | None |
| **Hepatic NAG**<br>**(D.O/mg tissue)** | 0.03224±0.0017 | 0.04614±0.003<sup>a,c</sup> | 0.03174±0.0053 | a: p= 0.0421 vs Sham<br>c: p= 0.0351 vs OVXR |
| **Adipocytes area**<br>**(mm<sup>2</sup>)** | 171.5±15.33 | 411.7±46.23<sup>a,c</sup> | 271.7±26.31 | a: p<0.0001 vs Sham<br>c: p=0.0054 vs OVXR |

SHAM: control group; OVX, ovariectomized; OVXR (OVX + caloric restriction); \*tissue weight (g)/ animal body weight X 100. Data express mean ± std error of mean (n=6/group) and values were compared by one-way ANOVA with Tukey post-test.

Serum cholesterol levels were lower in the OVXR group than in the OVX group (Figure 3A). Serum triglycerides (Figure 3B) and VLDL (Figure 3E) were similar among the groups. LDL-cholesterol (Figure 3D) was higher in the OVX group than in the Sham group. Normalization of the HDL-cholesterol fraction was observed for OVXR mice compared to Sham mice (Figure 3C), and HDL-cholesterol was lower in OVX mice than in Sham mice. Hepatic cholesterol levels were similar among the groups, but hepatic triglycerides were higher in OVXR mice than in Sham and OVX mice (Table 1). This result is mirrored by the liver tissue histology, which showed lipid vacuoles in the livers of OVXR mice (Figure 1F), suggesting metabolic adaptation to the CR diet. Similar adipocyte diameter (Table 1) was observed in the OVXR and Sham groups, whereas adipocyte diameter was increased in the OVX group when compared to that in the OVXR group.

### Oxidative stress balance

CAT protein expression was higher in OVX mice than in OVXR mice (Figure 4A), and was similar in OVXR and Sham mice. SOD-2 protein expression (Figure 4B) was similar among the groups. Representative bands of CAT, SOD-2, and beta-actin protein are shown in Figure 4C. Analysis of TBARS content in the serum (Figure 4D) and liver (Figure 4E) and H_2_O_2_ levels (Figure 4F) showed lower levels in the OVXR group than in the OVX group, suggesting reduced oxidative stress in the OVXR group compared to that in the OVX and Sham groups.

**Fig. 4.**
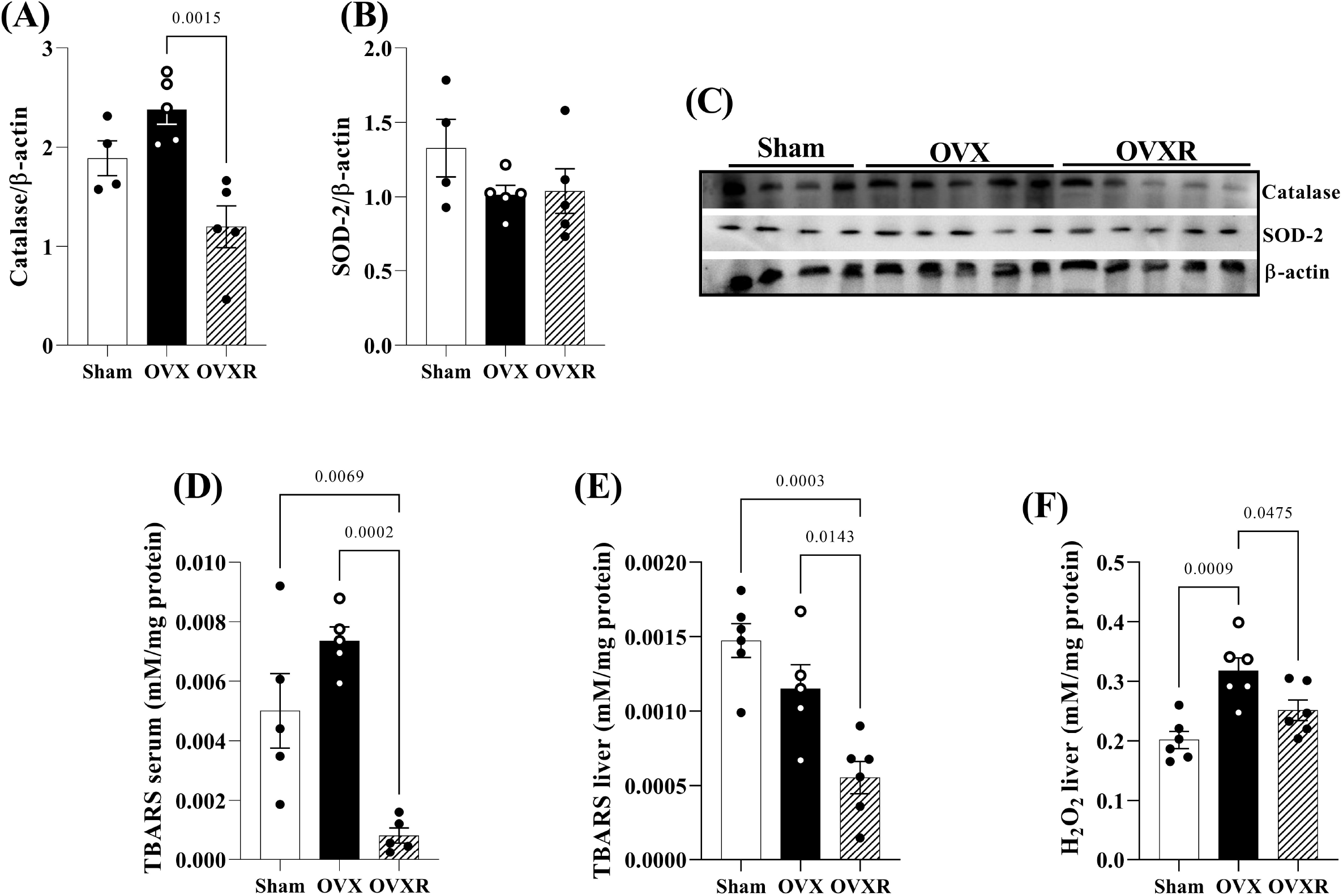
Typical immunoblots of (A) catalase (CAT), (B) superoxide dismutase (SOD2), and (C) representative blots. (D) Serum TBARS, (E) liver TBARS, and (F) liver H_2_O_2_ levels in the Sham, OVX, and OVXR groups. Data are mean ± SEM and were analyzed by one-way ANOVA followed by post-hoc Tukey test. The numbers of mice (n) and p-values are indicated on each graph.

### Hepatic inflammatory balance

Macrophage infiltration (NAG activity) in liver tissue was similar in the OVXR and Sham groups (Table 1) but was higher in the OVX group than in the Sham and OVXR groups. Levels of IL-6, a pro-inflammatory cytokine, were higher in OVX mice than in Sham mice (Figure 5A), whereas levels of IL-10, anti-inflammatory cytokine, were lower in OVX mice than in Sham mice (Figure 5B). Similar TGF protein expression was observed in all groups (Figure 5C).

**Fig. 5.**
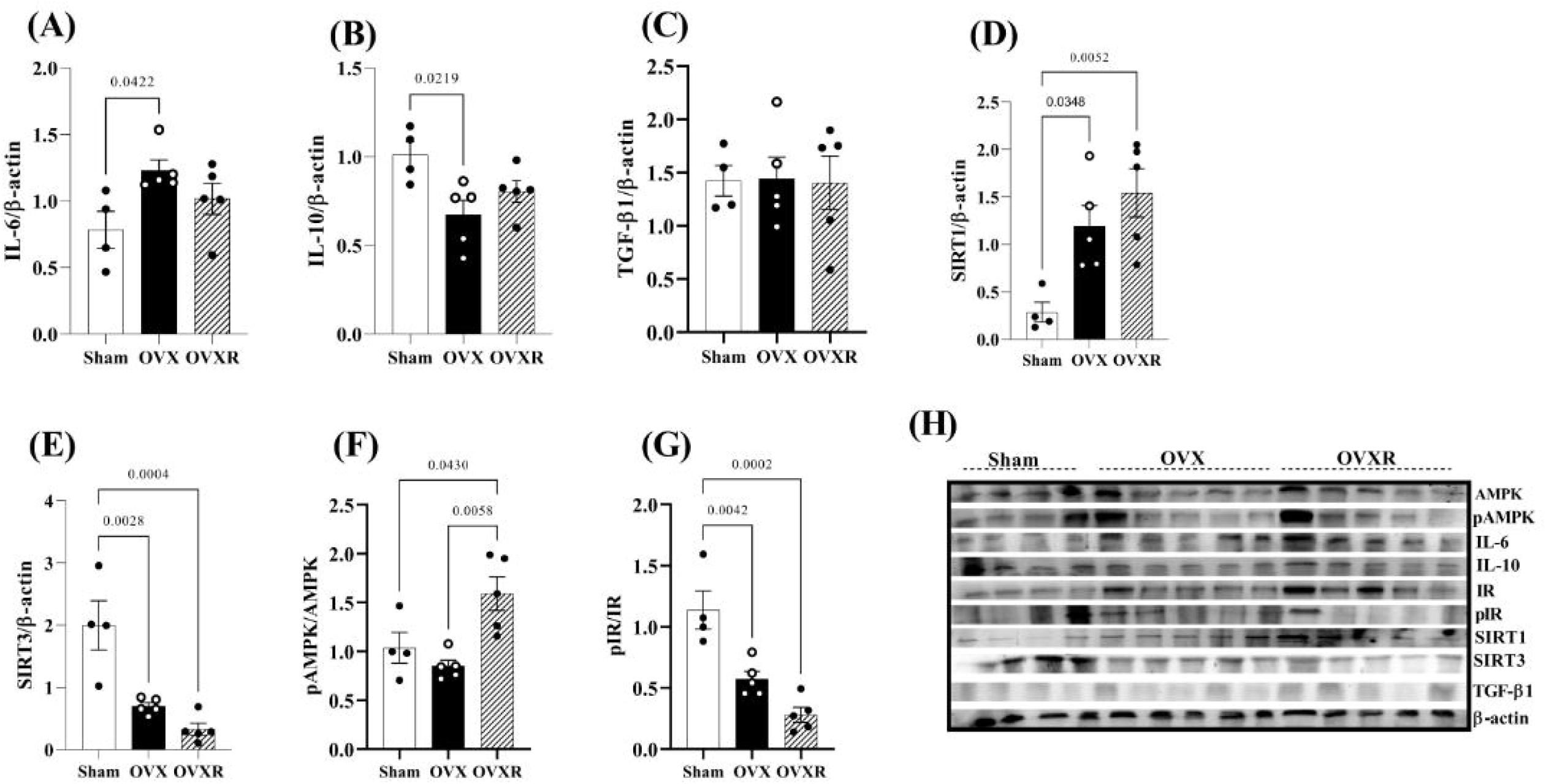
Typical immunoblots of (A) IL-6, (B) IL-10, (C) TGF-β1, (D) SIRT1, (E) SIRT3, (F) pAMPK/AMPK, (G) pIR/IR, and (H) representative blots in the livers of mice in the Sham, OVX, and OVXR groups. Data are mean ± SEM and were analyzed by one-way ANOVA followed by post-hoc Tukey test. The numbers of mice (n) and p-values are indicated on each graph.

### Hepatic metabolism

SIRT1 protein expression in the liver tissue was higher in OVXR mice than in Sham mice (Figure 5D). In contrast, SIRT3 (Figure 5E) and IR (Figure 5G) protein levels were lower in OVXR and OVX mice than in Sham mice. AMPK protein expression was higher in OVXR mice than in OVX and Sham mice (Figure 5F). Representative protein bands of AMPK, pAMPK, IL-6, IL-10, IR, pIR, SIRT1, SIRT3, TGF, and beta-actin are shown in Figure 5H.

## Discussion

CR is a well-accepted dietary intervention that can restore glucose homeostasis and promote weight loss [33]. In this study, we expected that CR would ameliorate the metabolic dysfunction and inflammation induced by ovariectomy. We observed weight loss and lower adiposity in ovariectomized mice under CR. These mice also had higher insulin sensitivity and normalized lipid profiles compared to the other mice. This can be attributed due a CR-induced homeostatic effect on oxidative stress and inflammatory balance in hepatic tissue.

Estrogen exerts protective effects on metabolism by regulating feeding behavior, glucose and lipid uptake, and the insulin pathway [34]. The reduction in estrogen level during menopause can lead to metabolism disruption, causing insulin resistance. In this study, CR decreased fasting glucose and insulin levels. Corroborating these findings, OVXR mice presented higher insulin sensitivity and glucose tolerance as evidenced by the ipGTT and ipITT results and the HOMA index. These beneficial changes in glycemic homeostasis can be attributed to increased expression of pAMPK/AMPK, but not pIR/IR, in the liver. AMPK controls glycemia and gluconeogenesis and is an important treatment target in diabetes [35]. Although the cholesterol fractions and triglycerides were unchanged, CR reduced total serum cholesterol.

In previous studies, ovariectomized mice displayed a nonalcoholic fatty liver disease (NAFLD) phenotype [36, 37]. Although we did not observe any changes in the cholesterol and triglyceride contents in the livers of OVX mice, triglycerides were increased in the liver of OVXR animals. Activation of AMPK inhibits hepatic lipogenesis by inactivating acetyl CoA carboxylase [38], resulting in fat oxidation [39]. Interestingly, AMPK expression induced hepatic lipid accumulation and fatty acid oxidation [35]. In this study, hepatic pAMPK/AMPK expression was significantly increased in OVXR mice, suggesting its involvement in hepatic lipid metabolism. This hepatic lipid storage in OVXR mice may be a transient metabolic process related to the restricted diet. Under CR, the OVXR mice used fat as an energy source to maintain cellular metabolism, which may explain the lipid accumulation in the liver. The mechanism may involve rapid response of AMPK phosphorylation promoting uptake of fatty acids stored in the liver [39, 35]. Our histological observations showed mild hepatic steatosis and a reduced adipocyte area following CR, corroborating the proposed mechanism. In a previous study, CR animals presented increased serum concentrations of free fatty acids and SIRT1, which may indicate mobilization of triglycerides by SIRT1 under CR [40].

Lipid accumulation increases oxidative stress and inflammation. This can exacerbate release of reactive oxygen species and pro-inflammatory cytokines, such as tumor-necrosis factor (TNF)-α and IL-6, in liver and adipose tissue. Previous studies have reported an increase in IL-6 in menopause [41, 42]. After menopause, levels of the anti-inflammatory cytokines IL-4, IL-10, and IL-12 are increased, which is related to an increase in TNF-α, as a compensatory mechanism [43]. In our study, OVX mice showed inflammatory effects, evidenced by an increase in IL-6 and a reduction in IL-10, due to the reduction in estrogen levels. The levels of these cytokines were unchanged in OVXR mice. Although TGF is an inflammatory factor that induces fibrosis, its expression was unaffected by ovariectomy. NAG is an indicator of macrophage activity and inflammation. The reduction in NAG level under CR suggests the anti-inflammatory effect of CR. NAG concentrations have been linked to obesity and diabetes [44].

In NAFLD, SIRT1 activation plays a beneficial role by decreasing oxidative stress and inducing deacetylation of PGC-1α, leading to reduced pro-inflammatory cytokine levels. Studies have shown that ovariectomy induced unfavorable cardiovascular changes and that estrogen treatment activated SIRT1 and promoted cardiovascular protection [45]. In mice, ovariectomy and SIRT3 depletion impaired the antioxidative system response and increased lipid damage, body weight gain, and the levels of oxidative stress-inducing genes [46]. Our findings showed that ovariectomy increased SIRT1 and decreased SIRT3, suggesting these proteins function in a regulatory mechanism to maintain normal metabolism.

The reductions in serum and liver TBARS contents and liver H_2_O_2_ levels in the OVXR mice suggest changes in the redox state of the liver as a protective mechanism against oxidative damage. The literature supports an increase in TBARS content following ovariectomy, corroborating the findings of the present study [47]. The reduction in TBARS contents in the OVXR group may explain the reduction in CAT and the similar SOD expression levels in the present study, suggesting a balance of mechanisms to combat oxidative stress under hypoestrogenism. The activity levels of the antioxidant enzymes SOD, CAT, and GPX were significantly diminished in obesity [48], and oxidative stress was enhanced as body weight increased in post-menopausal women [49]. Here, we aimed to understand the specific functions of CR in the “metabolic network” of tissues involved in whole-body glucose and lipid metabolism in ovariectomized mice. Notably, while CR benefitted the ovariectomized mice by reducing adiposity and increasing insulin sensitivity and glucose tolerance, suggesting a balance in oxidative stress, it induced negative changes in lipid metabolism in the liver, which was associated with changes in the protein expression of SIRT1 and SIRT3. Finally, the results of our study suggest that ovariectomy can accelerate metabolic changes from tissue damage to the body and that CR was able to restore, at least in part, glycemic homeostasis.

## Author statements

## Acknowledgments

We thank Ana Cristina Pires Menegheti, Bruno Alves Cia, Lucas Orzari, Mateus Eduardo Bortolanza da Silva, and Renata Barbieri for technical assistance.

## Author contribution statement

**Sapatini LRL**: Data curation, Investigation, Visualization, Writing – original draft. **Calsa B:** Data curation, Formal analysis, Investigation, Validation, Visualization, Writing – original draft, review and editing. **Marim LJ:** Investigation. **Helaehil JV:** Data curation, Investigation. **Amaral MEC:** Conceptualization, Data curation, Formal analysis, Funding acquisition, Investigation, Project administration, Resources, Supervision, Validation, Visualization, Writing – review & editing.

## Data availability

Data generated or analyzed during this study are available form the corresponding author upon reasonable request.

## Preprint or repository availability

bioRxiv preprint doi: https://doi.org/10.1101/2023.02.09.527836

